# High expression of CD38 and MHC class II on CD8^+^ T cells during severe influenza disease reflects bystander activation and trogocytosis

**DOI:** 10.1101/2021.02.09.430410

**Authors:** Xiaoxiao Jia, Brendon Y Chua, Liyen Loh, Marios Koutsakos, Lukasz Kedzierski, Moshe Olshanski, William R Heath, Jianqing Xu, Zhongfang Wang, Katherine Kedzierska

## Abstract

Although co-expression of CD38 and HLA-DR on CD8^+^ T cells reflects activation during influenza, SARS-CoV-2, Dengue, Ebola and HIV-1 viral infections, high and prolonged CD38^+^HLA-DR^+^ expression can be associated with severe and fatal disease outcomes. As the expression of CD38^+^HLA-DR^+^ is poorly understood, we used mouse models of influenza A/H7N9, A/H3N2 and A/H1N1 infection to investigate the mechanisms underpinning CD38^+^MHC-II^+^ phenotype on CD8^+^ T-cells. Our analysis of influenza-specific immunodominant D^b^NP_366_+CD8^+^ T-cell responses showed that CD38^+^MHC-II^+^ co-expression was detected on both virus-specific and bystander CD8^+^ T-cells, with increased numbers of both CD38^+^MHC-II^+^CD8^+^ T-cell populations observed in the respiratory tract during severe infection. To understand the mechanisms underlying CD38 and MHC-II expression, we also used adoptively-transferred transgenic OT-I CD8^+^ T-cells recognising the ovalbumin-derived K^b^SIINFEKL epitope and A/H1N1-SIINKEKL infection. Strikingly, we found that OT-I cells adoptively-transferred into MHC-II^−/−^ mice did not display MHC-II after influenza virus infection, suggesting that MHC-II was acquired via trogocytosis in wild-type mice. Additionally, detection of CD19 on CD38^+^MHC II^+^ OT-I cells further supports that MHC-II was acquired by trogocytosis, at least partially, sourced from B-cells. Our results also revealed that co-expression of CD38^+^MHC II^+^ on CD8^+^ T-cells was needed for the optimal recall ability following secondary viral challenge. Overall, our study provides evidence that both virus-specific and bystander CD38^+^MHC-II^+^ CD8^+^ T-cells are recruited to the site of infection during severe disease, and that MHC-II expression occurs via trogocytosis from antigen-presenting cells. Our findings also highlight the importance of the CD38^+^MHC II^+^ phenotype for CD8^+^ T-cell memory establishment and recall.

**Summary:** Co-expression of CD38 and MHC-II on CD8^+^ T cells is recognized as a classical hallmark of activation during viral infections. High and prolonged CD38^+^HLA-DR^+^ expression, however, can be associated with severe disease outcomes and the mechanisms are unclear. Using our established influenza wild-type and transgenic mouse models, we determined how disease severity affected the activation of influenza-specific CD38^+^MHC-II^+^CD8^+^ T cell responses *in vivo* and the antigenic determinants that drive their activation and expansion. Overall, our study provides evidence that both virus-specific and bystander CD38^+^MHC-II^+^ CD8^+^ T-cells are recruited to the site of infection during severe disease, and that MHC-II expression occurs, at least in part, via trogocytosis from antigen-presenting cells. Our findings also highlight the importance of the CD38^+^MHC II^+^ phenotype for CD8^+^ T-cell memory establishment and recall.

## Introduction

Influenza virus infections lead to annual seasonal epidemics that result in 243,000-640,000 deaths globally every year[1]. In 2017 alone, ~9.5 million people were hospitalized with lower respiratory tract influenza infections for a total of 81.5 million hospital days[2]. Furthermore, disease severity and mortality rapidly escalate when a new influenza A virus (IAV) emerges, for which a matching vaccine is unavailable, as exemplified by the catastrophic 1918-1919 H1N1 pandemic and the emergence of avian-derived H5N1 and H7N9 strains in 2003[3] and 2013[4], respectively. In the absence of neutralizing antibodies, the recall of pre-existing broadly cross-reactive memory CD8^+^ T cells that recognize conserved peptide epitopes derived from internal viral proteins presented by infected cells, can promote recovery against novel influenza A viruses (IAVs) and even different influenza types[5],[6],[7]. Activated CD8^+^ T cells can kill virus-infected cells via perforin- and granzyme-mediated lysis but also produce cytokines including IFN-γ and TNF-α, which render the local environment non-permissive for virus replication and ultimately provide some degree of heterosubtypic protection in the antibody-naïve individuals.

The importance of broadly cross-reactive CD8^+^ T cell responses in mitigating influenza disease severity has been shown during outbreaks caused by the emergence of novel subtypes, including the 2003 H5N1[8], 2009 pandemic H1N1[7] and 2013 H7N9 strains[9,10]. Our analysis of immune responses in H7N9-infected hospitalized patients in China demonstrated that early proliferation of H7N9-specific CD8^+^ T cells contribute to disease resolution and survival, as reflected by mild to moderate clinical scores and early time to discharge from hospital[9]. In particular, we showed that early, transient co-expression of CD38^+^ and HLA-DR^+^ (MHC-II^+^) on CD8^+^ T cells during the early stages of H7N9 infection was strongly associated with patient survival[10].

Co-expression of CD38, a cyclic ADP ribose hydrolase and cell surface glycoprotein involved in signal transduction, adhesion and calcium signalling[11], and MHC-II on CD8^+^ T cells is recognized as a classical hallmark of activation during viral infections. CD38^+^MHC-II^+^CD8^+^ T cells can be detected transiently at high levels in patients infected with viruses such as Ebola[12], HCV[13], HIV [14],[15], Dengue[16], pandemic H1N1 IAV[17], and more recently SARS-CoV-2[18],[19] during the acute phase of infection. CD38^+^MHCII^+^CD8^+^ T cells are known for their proliferative capacity, exhibit effector functions such as cytotoxicity and cytokine production[20],[21],[22] and typically decrease in frequency following the resolution of infection[21,22],[18]. However, in converse prolonged expression and high frequency of CD38^+^MHCII^+^CD8^+^ T cells can be associated with a loss of effector function, as shown during chronic viral infections[23,24]. Moreover, H7N9-infected patients with fatal disease outcomes displayed high and prolonged expression of CD38^+^MHC-II^+^ on CD8^+^ T cells, which had minimal capacity to produce IFN-γ, thus linking the prominence of CD38^+^MHC-II^+^ phenotype to fatal outcomes in severe IAV-mediated pneumonia[10].

Despite the important role of CD38^+^MHCII^+^CD8^+^ T cells in viral control, the underlying mechanisms driving their activation and regulation during viral infections are still poorly understood. Here, using our established H7N9 and H3N2 influenza wild-type and transgenic mouse models, we determined how disease severity affected the activation of IAV-specific CD38^+^MHC-II^+^CD8^+^ T cell responses *in vivo* and the antigenic determinants that drive their activation and expansion. Our study also revealed how this population acquires MHC-II expression from antigen-presenting cells by trogocytosis and revealed superior memory persistence of CD38^+^MHC-II^+^CD8^+^ T cells.

## Results

### H7N9 infection in mice induces CD38^+^MHC-II^+^PD-1^+^ activation profiles

Our previous data revealed that patients hospitalized with severe H7N9 influenza had high and prolonged expression of CD38, MHC-II and PD-1 on CD8^+^ T cells[10]. To understand the mechanisms underlying such hyper-activated CD38, MHC-II and PD-1 phenotype, we first investigated the recruitment kinetics and specificity of CD38^+^MHC-II^+^ and CD38^+^PD-1^+^ CD8^+^ T cells in a mouse model of H7N9-infection in C57BL/6 (B6) mice. To minimize H7N9-induced mortality, B6 mice were first primed intranasally (i.n.) with 10^4^ EID50 of the avirulent strain A/Chicken/Shanghai/F/98 (H9N2) 60 days prior to i.n. challenge with 10^6^ pfu of A/Shanghai/4664T/2013 (H7N9) virus (Fig 1A). Overall, ~20% of CD8^+^ T cells in the spleen or mediastinal lymph node (MLN) were CD38^+^MHC-II^+^ (20.2% for both spleen and MLN) (Fig.1C i,ii, Fig.S1) or CD38^+^PD-1^+^ (range 17.8-20.6%) (Fig.1C ii,iii) on d10 after H7N9 exposure. Analysis of CD8^+^ T-cell responses directed at the immunodominant D^b^NP_366_ epitope showed 3-5-fold enrichment of D^b^NP_366_^+^CD8^+^ T cells within the CD38^+^MHC-II^+^ or CD38^+^PD-1^+^ sets from the spleen and MLN (Fig 1B, Fig S1). Alternatively, when gated on the D^b^NP_366_^+^CD8^+^ set, the CD38^+^MHC-II^+^ (59.5% and 81.1% for spleen and MLN) and CD38^+^PD-1^+^ (90.2% and 93.4%) phenotypes were prominent (Fig 1C), with the CD38 and MHC-II levels (measured by frequency) being higher for MLN than spleen (p<0.05) (Fig 1C). Such high expression of CD38^+^MHC-II^+^ or CD38^+^PD-1^+^ on tetramer-specific CD8^+^ T cells after H7N9 infection in mice supports the observations from H7N9 patients. As expected, the majority of CD38^+^MHC-II^+^ or D^b^NP_366_^+^CD8^+^ T cells in MLN and spleen were CD44^hi^ (antigen-experienced), though expression of CD62L, a molecule that mediates trafficking into lymph nodes via high endothelial venules, differed for these two sites (Fig 1D).

**Figure 1.**
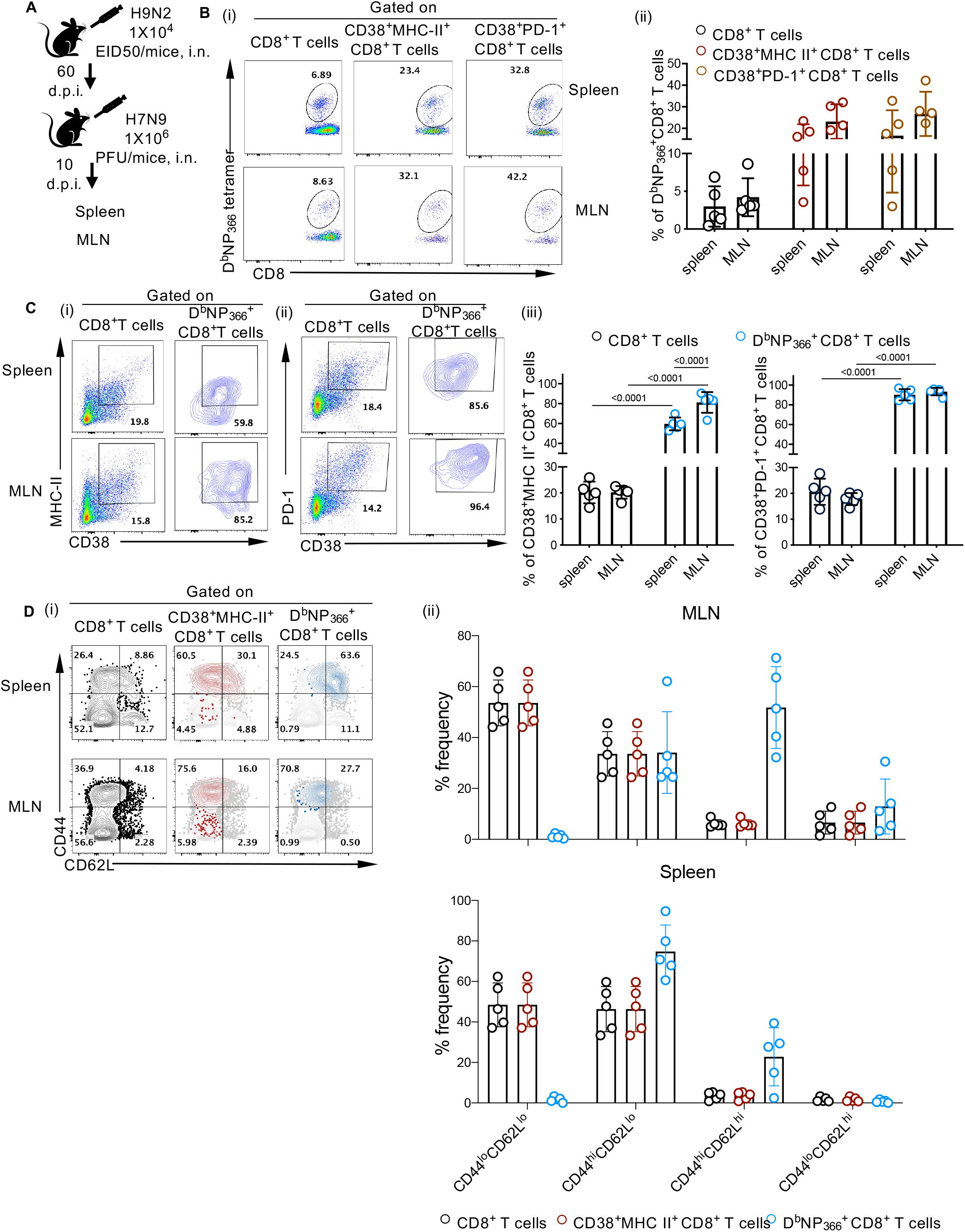
H7N9 infection in mice induces CD38^+^MHC-II^+^PD-1^+^ activation profiles. **(a)** C57BL/6 Mice (n=5) were infected i.n. with H9N2 (A/Chicken/Shanghai/F/98) (10^4^ EID50) virus and challenged i.n. at 60 d.p.i.with H7N9 (A/Shanghai/4664T/2013) (10^6^ PFU) virus. Spleens and MLNs were harvested 10 days after H7N9 challenge and phenotype of CD8^+^ T cells analysed. **(b)** (i) Representative dot plots and (ii) frequencies of D^b^NP_366_^+^CD8^+^ T cells within CD8^+^, CD38^+^MHC-II^+^CD8^+^ and CD38^+^PD-1^+^CD8^+^ gated populations are shown. **(c)** (i) Depicted are representative dot plots and (ii) frequencies of CD38^+^MHC-II^+^ or CD38^+^PD-1^+^ within CD8^+^ and D^b^NP_366_^+^CD8^+^ T cell populations. (d) Memory profiles were also analysed based on co-expression of CD62L and CD44. Statistics were performed using Mann–Whitney test.

As CD8^+^ T cells from H7N9-infected hospitalized patients displayed high PD-1 levels, we compared PD-1 expression on H7N9-specific D^b^NP_366_^+^CD8^+^ T cells to that on CD38^+^MHC-II^+^ or CD38^−^MHC-II^−^ CD8^+^ T cells. D^b^NP_366_^+^CD8^+^ T cells in both the MLN and spleen displayed the highest level of PD-**1 (Fig 2).** As expected, PD-1 levels on CD8^+^ T cells negative for dual expression of CD38^+^MHC-II^+^ were minimal. Thus, high CD38^+^MHC-II^+^PD-1^+^ expression on D^b^NP_366_^+^CD8^+^ and CD38^+^MHC-II^+^ CD8^+^ T cells confirmed the correlation between high CD38^+^MHC-II^+^PD-1^+^ CD8^+^ T-cell prevalence and severe disease.

**Figure 2.**
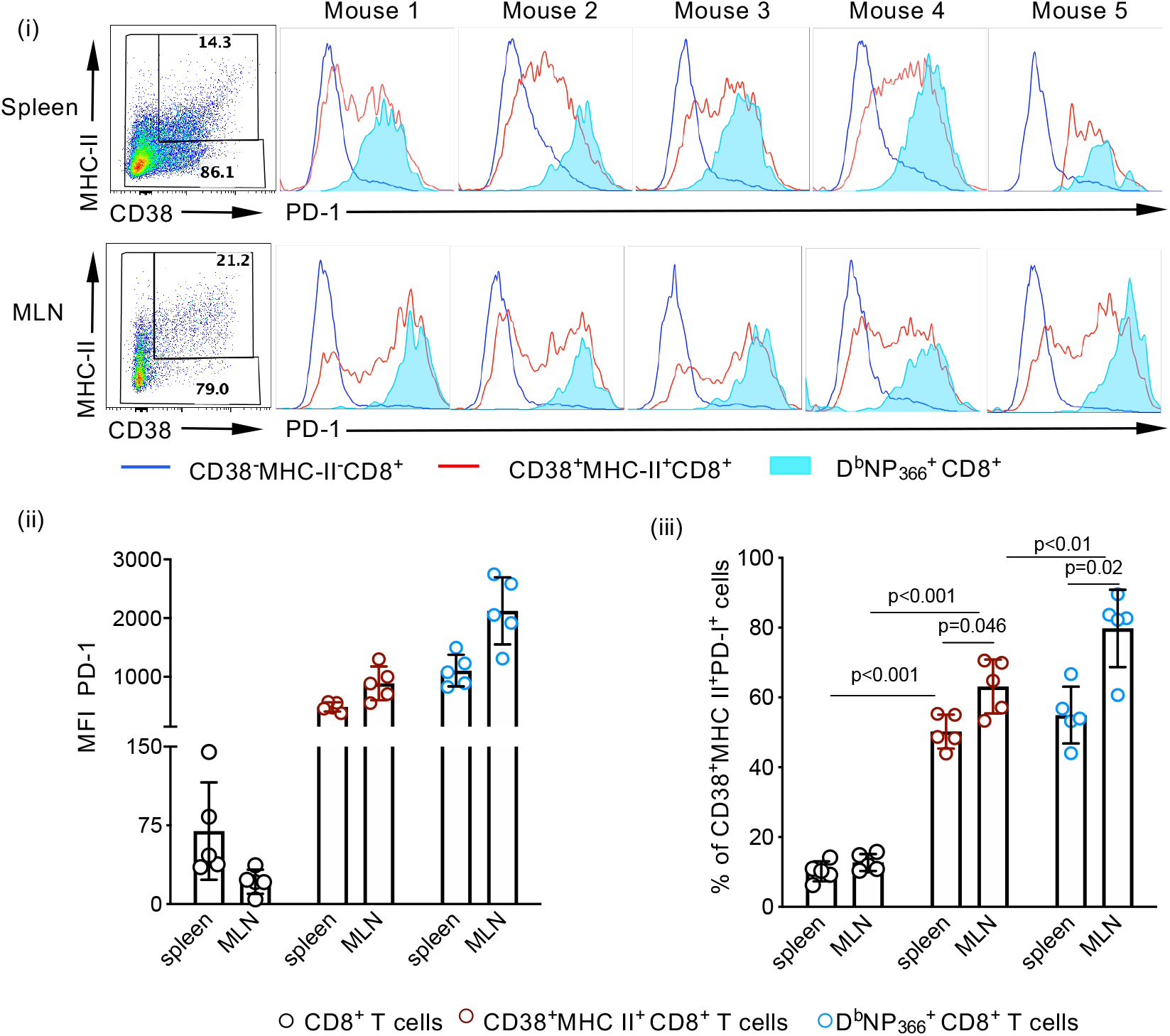
CD38^+^MHC-II^+^CD8^+^ T cells express higher levels of PD-1. Spleens and MLNs of C57BL/6 mice (n=5) infected i.n. with H9N2 (10^4^ EID50) and again 60 days later with H7N9 (10^6^ PFU) were obtained 10 days following the last challenge. (a) The expression of PD-1 on D^b^NP_366_^+^CD8^+^, CD38^+^MHC-II^+^CD8^+^ T cells and CD38^−^MHC-II^−^ CD8^+^ T cell populations from each animal is depicted as (i) histograms, (ii) mean fluorescence intensity (MFI) and (iii) frequency. Statistics were performed using Mann–Whitney test.

### Recruitment of CD38^+^MHC-II^+^ and CD38^+^PD-1^+^CD8^+^ T cells to the site of infection

To determine how virus load and disease severity affect the recruitment kinetics and phenotype of CD38^+^MHC-II^+^ and CD38^+^PD-1^+^ CD8^+^ T cells, we infected B6 mice intranasally with a low (10^2^ pfu) or high (10^5^ pfu) dose of A/HK/x31 (H3N2; X31) virus (Fig 3A). While comparable virus replication kinetics were observed at d3, d5, d7 and d10 after infection (Fig 3B), the high dose caused more severe disease, as evidenced by significantly greater body weight loss between d3 and d7 (p<0.001) (Fig 3C), and increased lethality (p=0.003) (Fig 3D) as well as excessive inflammation in the lung (Fig 3E). The 10^5^ pfu X31 virus challenge was associated with more rapid acquisition of a PD-1^hi^ phenotype (p<0.05 on d7) on D^b^NP_366_^+^CD8^+^ T cells (Fig. 3F), relecting the pattern found previously for H7N9-infected mice and patients who succumbed to H7N9 disease.

**Figure 3.**
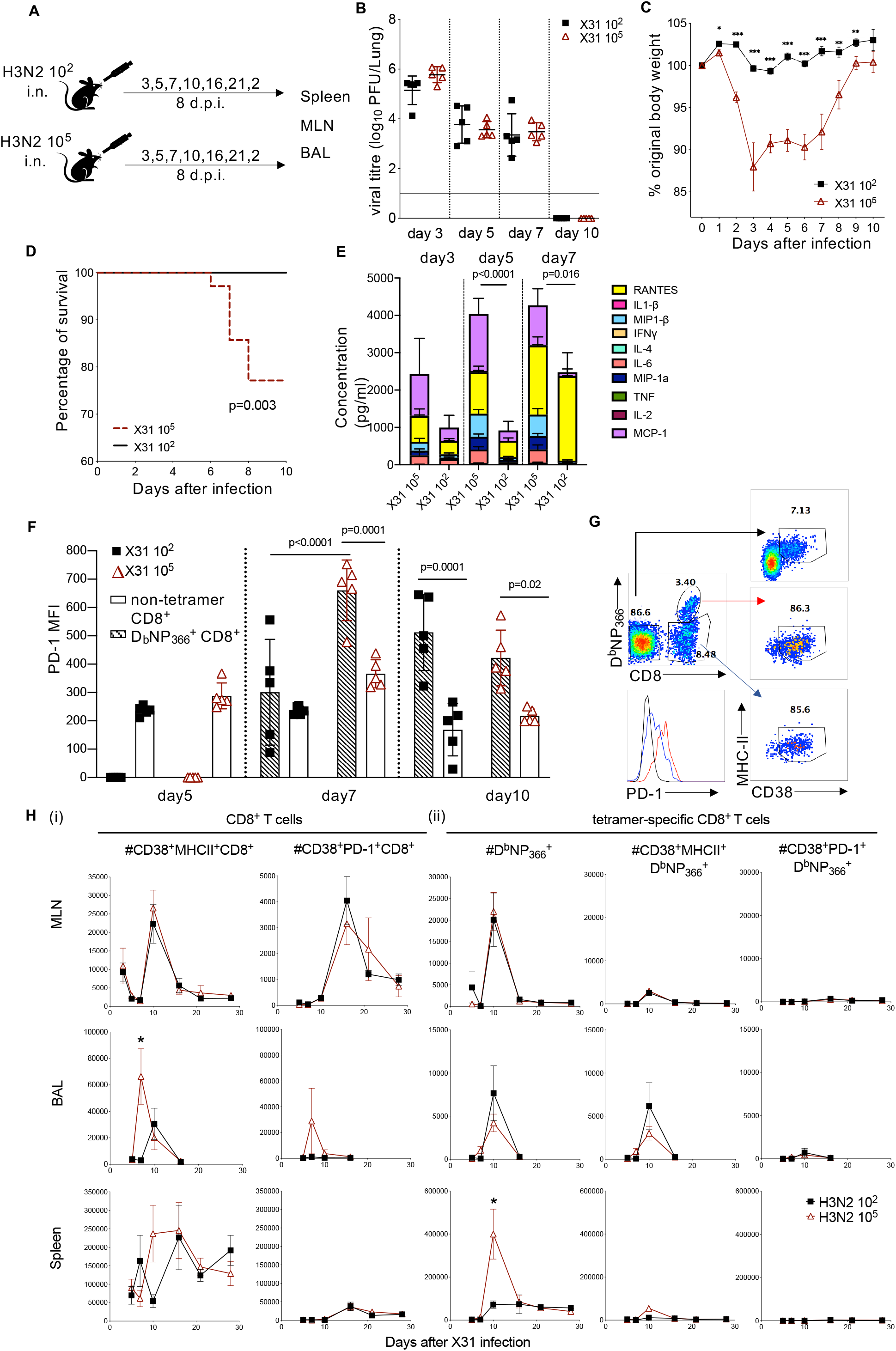
H3N2 viral dose affects kinetics of CD38^+^MHC-II^+^ and CD38^+^MHC-II^+^PD-1^+^ at the site of infection. **(a)** C57BL/6 mice were infected i.n. with X31 at either a high (10^5^ PFU) or low (10^2^ PFU) viral dose (n=5/group). **(b)** At various days post-infection (p.i.), lung viral titres were determined in MDCK plaque assays. Each symbols denotes an individual mice and the mean (±SD) is shown for each group. Mice were also monitored daily after infection for **(c)** weight loss and **(d)** survival (mice were humanely culled when a weight loss of >20% of their pre-experimental body weight was observed over a 2 day period and accompanied by clinical signs of disease, or when a weight loss of >25% of their pre-experimental body weight occurred). **(e)** Lung homogenates were assayed by cytometric bead array to determine the cumulative amount of total cytokines present and the amounts of each cytokine. **(f)** MFI of PD-1 on D^b^NP_366_^+^CD8^+^ T cells (dashed bars) or non-tetramer CD8^+^ T cells (open bars). **(g)** Representative histograms of PD-1 expression on D^b^NP_366_^+^ (red line) or D^b^NP ^−^specific (blue line) CD38^+^MHC-II^+^CD8^+^ T cells or remaining D^b^NP ^−^CD8^−^ T cells (black line). **(h)** Numbers of (i) non-tetramer- and (ii) D^b^NP366-tetramer specific CD38^+^MHC-II^+^ and CD38^+^MHC-II^+^PD-1^+^ CD8^+^ T cells in MLN, BAL and spleen at various time-points after infection. *p<0.05, **p<0.01 using Mann–Whitney test.

Although cell numbers for all CD38^+^MHC-II^+^ and CD38^+^PD-1^+^ CD8^+^ T cells in the regional MLN were comparable from d5 and beyond (Fig 3H (i)), only a small population of cells (11.8%) were D^b^NP_366_^+^ (Fig 3H (ii)) within CD38^+^MHC-II^+^ CD8^+^ T cells (peak d10) and (17.3%) within CD38^+^PD-1^+^CD8^+^ T cells (peak d16), indicating that the majority of CD38^+^MHC-II^+^ and CD38^+^PD-1^+^ CD8^+^ T cells recruited during severe influenza disease were tetramer-negative. Furthermore, CD38^+^MHC-II^+^ and CD38^+^PD-1^+^ CD8^+^ T cells were increased and peaked earlier (d7) at the site of infection (BAL) following the high dose challenge (Fig 3H i left, Fig S2), while the IAV-specific CD8^+^ T cell responses peaked 3 days later (Fig 3H ii left, Fig S2) and at comparable numbers for both (10^5^ and 10^2^ pfu) groups. Together, these data suggest that D^b^NP366 tetramer-negative CD38^+^MHC-II^+^CD8^+^ T-cells, many of which numerically will not be virus-specific, extravasate into the lung prior to recruitment of the majority of antigen-specific cells. Whether this is a consequence or a cause of more severe inflammation and pathology is currently unclear.

### CD38^+^CD8^+^ T cells utilise trogocytosis to acquire expression of MHC-II

As expression of the *H2-Ab (MHC-II)* gene at the transcriptional level depends on the transcription factor *CIITA*[25][26], and murine CD8^+^ T cells lack intrinsically expressed *CIITA*[27], we further investigated how murine CD8^+^ T cells acquire expression of the MHC-II. We utilised a transgenic OT-I mouse model in the influenza setting. OT-I CD8^+^ T cells contain a TCRαβ specific for the K^b^OVA_257-264_ (OVA; SIINFEKL) epitope, which proliferate after infection with influenza-OVA virus (with SIINFEKL engineered into the neuraminidase stalk)^53^. We adoptively transferred naïve OT-I CD8^+^ T cells into MHC II^−/−^ or C57BL/6 wild-type mice prior to X31-OVA infection (Fig 4A). We observed greater body weight loss (5%) in MHC II^−/−^ mice compared to wild-type mice particularly at 3-5 days post-infection (Fig 4B), indicating that lack of MHC-II expression leads to perturbed immune responses and increased disease severity. When we analysed the expansion of OT-I cells (Fig 4Ci) by frequency (Fig 4Cii) or number (Fig 4Ciii) of the total CD8^+^ T cell pool, we observed similar numbers in BAL, MLNs and spleens of both WT and MHC II^−/−^ mice. As there were also no differences in the combined total number of OT-I cells from these organs of each mouse (Fig 4C iv), this indicates that OT-I cells proliferate to the same extent in both WT and MHC−/− mice.

**Figure 4.**
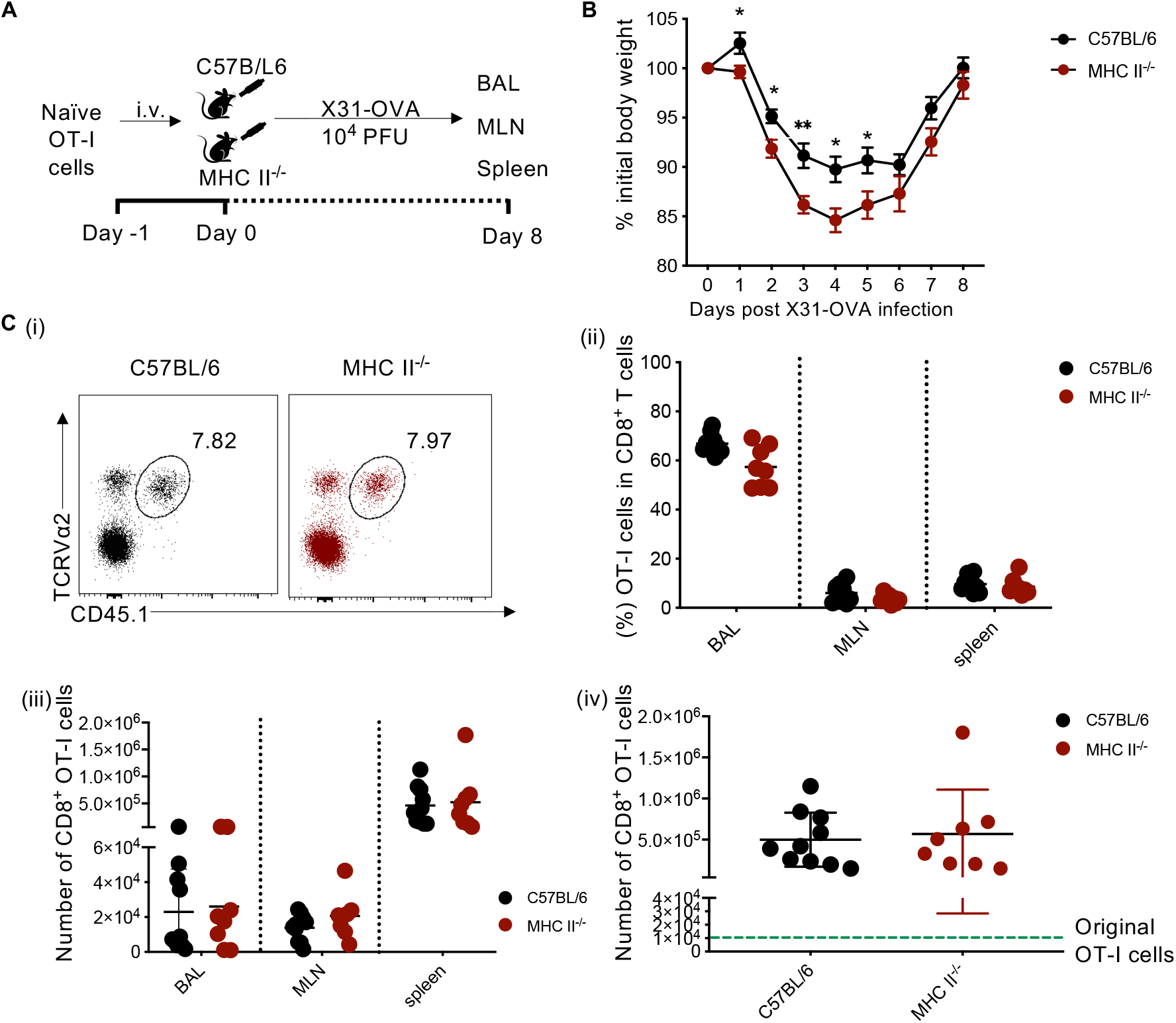
Similar OT-I T cell proliferation in C57BL/6 and MHC-II^−/−^ mice after IAV infection. **(a)** C57BL/6 and MHC-II^−/−^ mice (n=5/group) received 10^4^ OT-I T cells via the i.v. route 24 hrs prior to infection. Mice were subsequently infected with 10^4^ PFU of X31-OVA (H3N2) and BAL, MLN and spleen were harvested at 8 d.p.i.. **(b)** Mice were monitored daily during the course of infection for weight loss before **(c)** OT-I T cell populations each organ was enumerated. (i) Depicted are representative dot plots of CD45^+^Vα2^+^ OT-I T cells within the CD8^+^ T cell pool in the MLN. Shown are also (ii) percentages, (iii) numbers of OT-I T cells in each different organs and (iv) as a cumulative total. *p<0.05, **p<0.01.

Analysis of MHC-II expression on OT-I cells in BAL, MLNs and spleens of MHC-II^−/−^ or B6 mice showed that MHC-II phenotype was only limited to OT-I cells transferred into B6 mice and absent in those transferred into MHC-II^−/−^ mice (Fig 5Ai-iii). This observation extended to CD38^+^ OT-I cells, thus indicating that murine CD8^+^ T cells do not intrinsically express MHC-II but acquire it from other cells during influenza virus infection.

**Figure 5.**
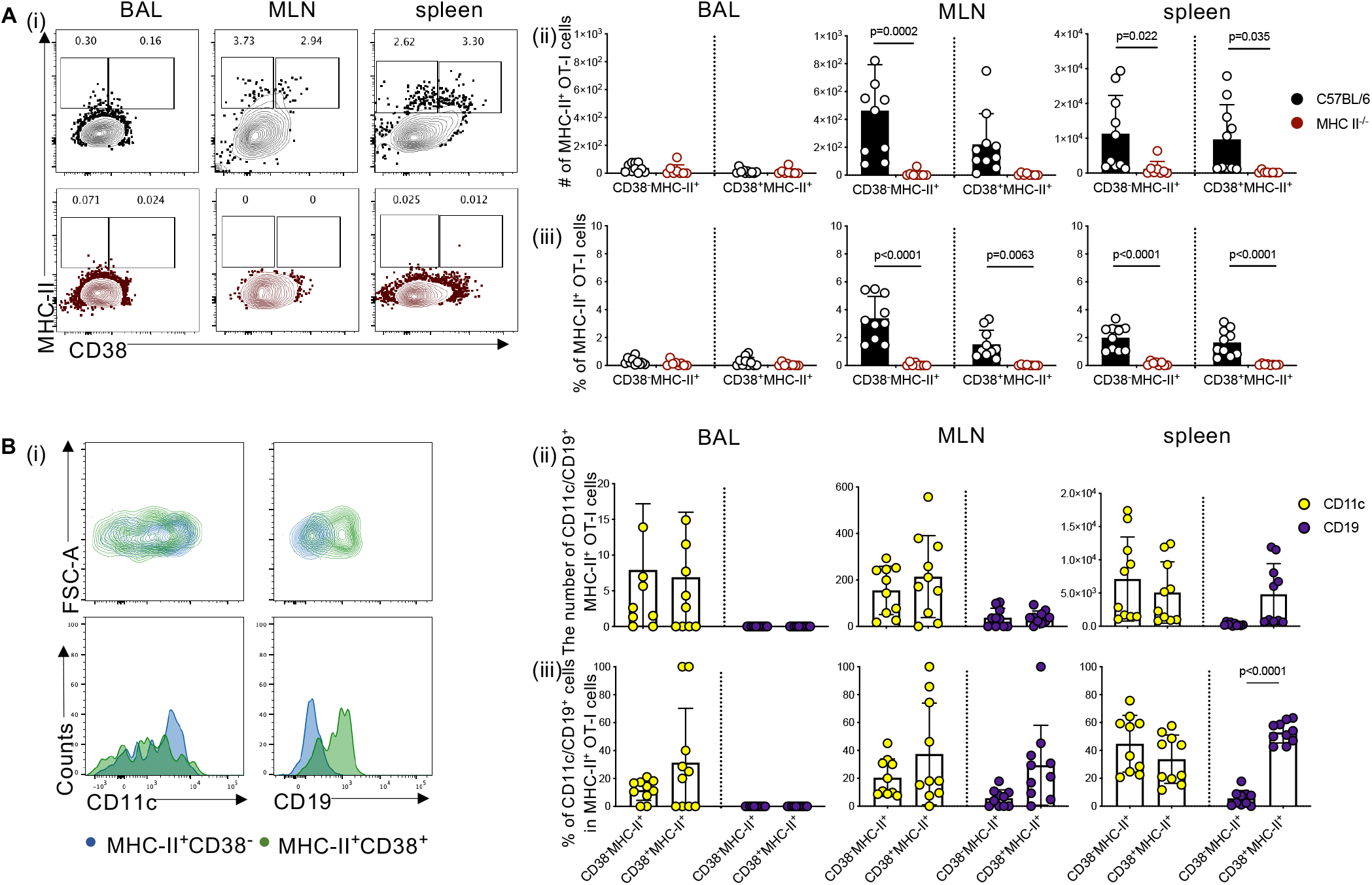
CD38^+^ OT-I T cells do not intrinsically express MHC-II and acquire it via trogocytosis. **(a)** C57BL/6 and MHC-II^−/−^ mice (n=5/group) received 10^4^ OT-I T cells via the i.v. route 24 hrs prior to infection with 10^4^ PFU of X31-OVA. (i) Representative dot plots of CD38 and MHC-II expression on OT-I T cells in MLN, BAL and spleens 8 day p.i. (ii) Bar graphs show the (ii) number and (iii) frequency of CD38^−^MHC-II^+^and CD38^+^MHC-II^+^ cells within the MHC-II^+^ OT-I T cell pool in each organ. **(b)** (i) Representative dot plots and histrograms of CD11c and CD19 expression on CD38^−^ MHC-II^+^ (green) and CD38^+^MHC-II^+^ (orange) OT-I T cells in C57BL/6 mice as well as (ii) their numbers and (iii) frequencies.

To determine how transferred OT-I cells could have acquired MHC-II, we stained OT-I cells in influenza-infected mice for markers associated with antigen presentation, including CD19 to represent B cells and CD11c as a surrogate DC and macrophage marker[28]. In BAL, MLN and spleen, CD11c^+^ OT-I cells were consistently detected on both CD38^−^MHC-II^+^ and CD38^+^MHC-II^+^ populations in the BAL (mean 11.7% and 31.4%) as well as the MLN (mean 20.5% vs 37.5%) and spleen (average ~ 44.8% vs 33.7%) (Fig 5Bi,ii and iii). In contrast, CD19 was associated mostly with CD38^+^MHC-II^+^ OT-I cells (mean 5.7% vs 29.4% in MLN and 5.6% vs 53.0% in spleen), while was rarely detected on CD38^−^MHC-II^+^ and no CD19 expression was observed on OT-I cells in BAL, suggesting that interaction of CD38^+^MHC-II^+^CD8^+^ T cells with CD19^+^ B cells via trogocytosis provides the source of MHC-II.

To further verify whether CD19, CD11c and MHC-II expression was intrinsic to OT-I cells, we measured CD19, CD11c and MHC-II (H2-Ab) mRNA levels in naïve, effector (8 d.p.i) and memory (30 d.p.i) CD8^+^ T cell populations following X31-OVA infection. Although gene expression of CD11c was not initially detected in naïve populations (Fig 6A; mean=14.67), a 2.4 ×10^2^-fold increase was observed in effector (Fig 6B; mean=3494) and 1.4 ×10^2^-fold increase in memory (Fig. 6C; mean=1985.5) CD8^+^ T cell populations, demonstrating that CD11c is intrinsic to CD8^+^ T cells and upregulated as they differentiate. In contrast, this was not the case for CD19 and MHC-II (H2-Ab), with undetectable levels of gene expression observed in naïve, effector and memory CD8^+^ T cell populations, similar to levels observed for CD4 gene expression. Overal, our data support the notion that the surface expression of MHC-II on CD8^+^ T cells is potentially acquired from B cells and dendritic cells/macrophages during influenza virus challenge.

**Figure 6.**
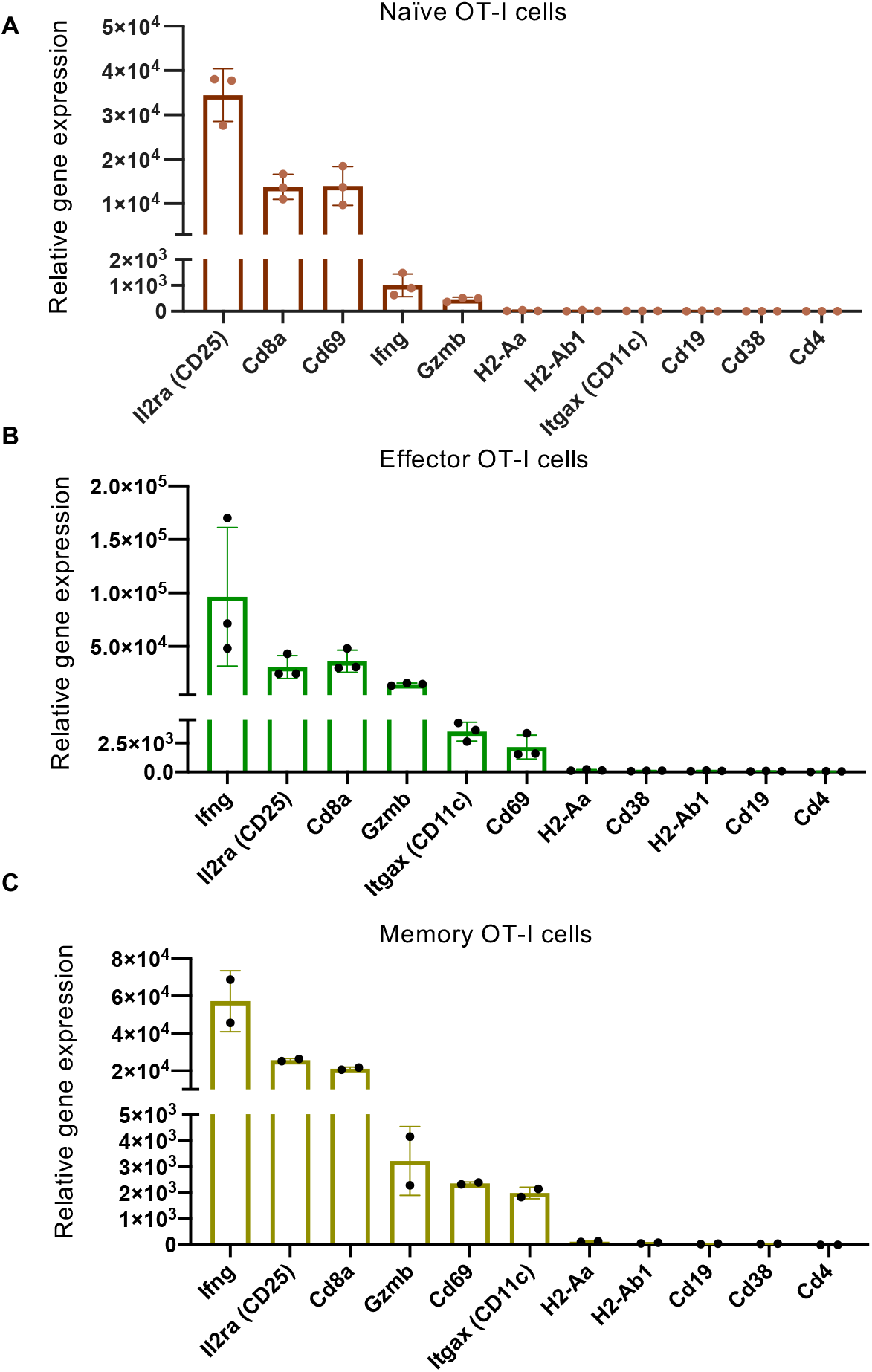
Gene expression in CD8^+^ T cells at naïve, effector and memory stage following influenza virus infection. Naïve OT-I cells were adoptively transferred into C57BL/6 mice and recipient mice were subsequently infected with 10^4^ PFU of A/HKx31-OVA. Immune organs were harvested at **(a)** 0 (naïve OT-1 cells), **(b)** 10 days (effector OT-I cells) or **(c)** 60 days (memory OT-I cells) post-infection, to analyse gene expression in CD8^+^ T cells by RNAseq. The original data were derived from Brendan E. Russ *et al*[55] and re-analysed for this study.

### Superior recall capacity of CD38^+^MHC-II^+^ CD8^+^ T cells

To investigate the functional significance of CD38^+^MHC II^+^CD8^+^ T cells during IAV infection, we analysed the recall capacity of memory OT-I cells established from 4 subsests of d8 effectors depicted by CD38 and MHC-II expression (CD38^−^MHC-II^−^, CD38^+^MHC-II^−^, CD38^−^MHC-II^+^, CD38^+^MHC-II^+^). OT-I cells adoptively transferred into B6 mice were expanded *in vivo* following infection with X31-OVA. On day 8 after infection, 4 populations of OT-I cells were FACS-isolated (Fig 7A). Each (CD38^−^MHC-II^−^, CD38^+^MHC-II^−^, CD38^−^MHC-II^+^, CD38^+^MHC-II^+^) population was adoptively transferred into separate naive receipient B6 mice, and subsequently rested for 30 days to establish memory formation. All mice were then challenged with X31-OVA virus. Mice that received CD38^−^MHC-II^−^ OT-I cells exhibited significantly (p<0.05) more body weight loss than mice that received OT-I cells expressing only CD38 (CD38^+^MHC II^−^), MHC II (CD38^−^MHC II^+^) or both these markers (CD38^+^MHC II^+^) (Fig 7B), indicating an important contribution of these activation markers for protection against severe IAV disease. Importantly, the greatest recall numbers of memory OT-I cells were observed in both the BAL (Fig 7C i,ii) and spleen (Fig 7C i,iii) of mice that received CD38^+^MHC-II^+^ cells with OT-I numbers being ~3.1-fold and ~4.2-fold higher, respectively, when compared to other subsets.

**Figure 7.**
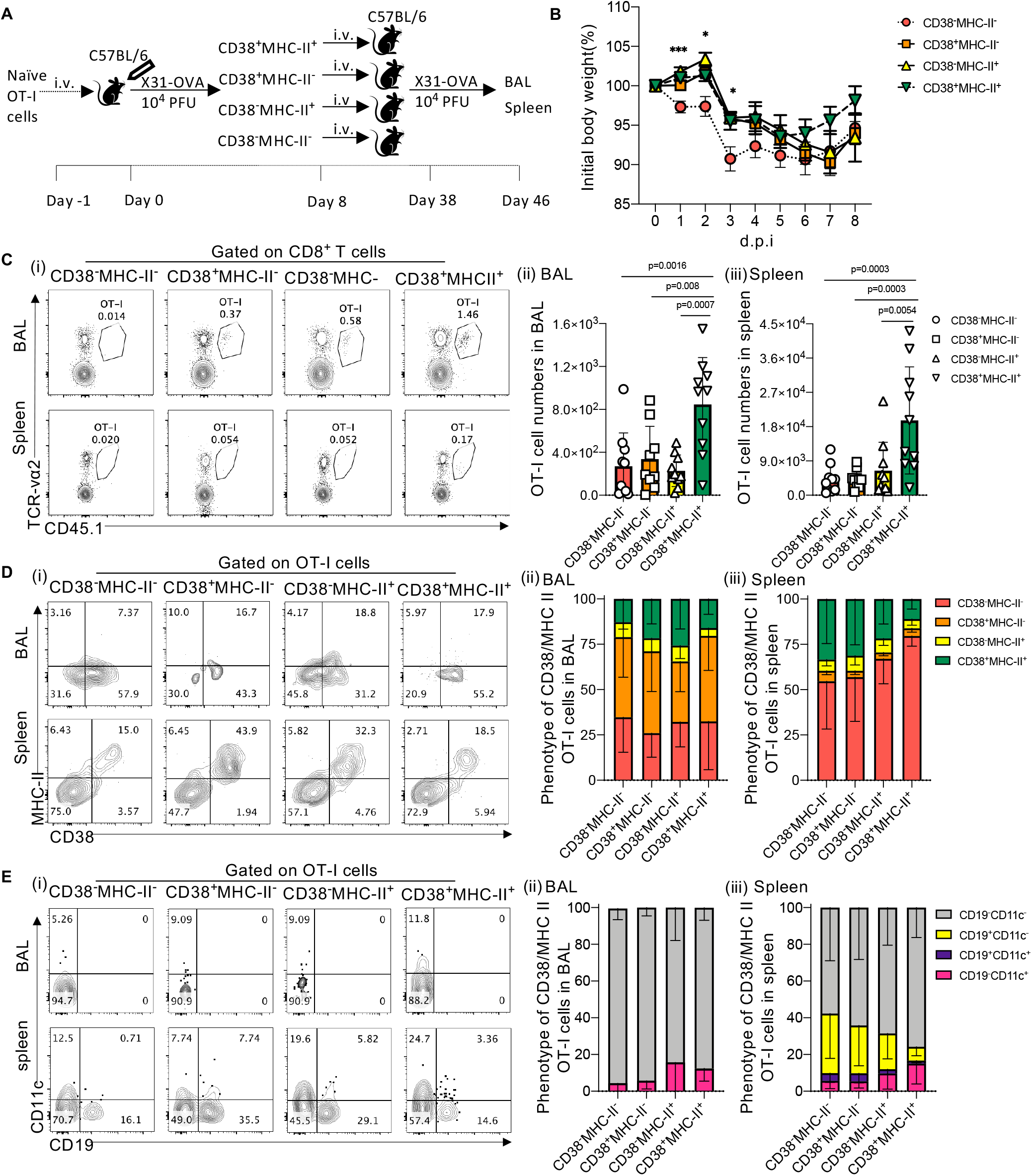
Superior recall capacity of CD38^+^MHC-II^+^ CD8^+^ T cells. **(a)** Naïve OT-I cells (10^4^ per mouse) were transferred into C57BL/6 mice and the next day challenged with 10^4^ PFU X31-OVA virus. OT-I cells (from MLN and spleen) were sorted based on CD38 and MHC-II expression on day 8 p.i. Sorted OT-I cells were adoptively transferred into new receipient C57BL/6 mice (3×10^5^ cells per mice) and rested for 30 days. Mice were then challenged with 10^4^ PFU X31-OVA virus and sacrificed on day 8 p.i. **(b)** Mice body weight were monitored over the course of 8 days following secondary viral challenge. **(c)** (i) Reprensentative FACS plots gated on CD8^+^ T cells across different receipient mice groups and showing the frequency of OT-I cells in BAL and spleen. Numbers of OT-I cells in (ii) BAL and (iii) spleen. **(d) (** i) Reprensentative FACS plot for CD38 and MHC-II expression on OT-I cells from different receipient mice groups. Percentage of phenotypes based on CD38 and MHC-II of OT-I cells in (ii) BAL and (iii) spleen. **(e)** (i) Reprensentive FACS plot for CD19 and CD11c expression on OT-I cells from different receipient mice groups. Percentage of phenotypes based on CD19 and CD11c of OT-I cells in (ii) BAL and (iii) spleen. *p<0.05, **p<0.01,

Given that we observed the acquisition of MHC-II molecules by CD8^+^ T cells during primary infection (Fig 5), we investigated whether this process also occurred following secondary challenge. Irrespective of the original phenotype of OT-I cells transferred, MHC-II^+^ OT-I cells (mean range from 20.4-31.06% across four groups) were found in BAL (Fig 7D i,ii). Furthermore, distinct populations of CD38^+^MHC-II^+^ OT-I cells were detected in the spleens of mice across all four groups (Fig 7D i,iii), indicating that OT-I cells that did not previously exhibit this phenotype can differentiate into CD38^+^MHC II^+^ expressing cells upon rechallenge. This, however, did not improve their recall ability and highlights the importance of establishing CD38^+^MHC-II^+^ phenotype prior to secondary exposure. Interestingly, while both CD19^+^ and CD11c^+^ OT-I cells could be found on CD38^+^MHC II^+^ OT-I cells in the spleens of all animal groups (Fig 7E i,iii), only CD19^−^CD11c^+^ OT-I cells could be detected in BAL (Fig 7E i,ii). Overall, our findings highlight the importance of the CD38^+^MHC II^+^ phenotype for CD8^+^ T-cell memory establishment and recall.

## Discussion

Co-expression of CD38^+^MHC-II^+^ on human CD8^+^ T cells is a hallmark of viral-specific T cell responses during many viral infections. In particular, early and rapid expansion of CD38^+^MHC-II^+^ CD8^+^ T cells can correlate with effective viremia control during acute HIV[20], Ebola[12], HCV[29], Dengue[16] infection. Our previous studies in H7N9-patients in China showed that the earlier recruitment of activated CD8^+^ T cells correlated with shorter duration of hospitalization, while late recruitment of CD8^+^ T cells leads to more diverse immune responses including upregulation of neutralizing antibodies, CD4^+^ T cells and NK cells[9]. While patients who recovered from H7N9 had early IFN-γ-secreting CD8^+^ T cells[10], persistent and prolonged expression of CD38 and MHC-II on CD8^+^ T cells was associated with fatal outcomes[30].

In this study, we demonstrate that IAV infection in a mouse model also induced the activation and expansion of CD38^+^MHC-II^+^ CD8^+^ T cells. Expression of CD38^+^ and MHC-II^+^ on CD8^+^ T cells recalled following H7N9 challenge correlated with high levels of PD-1 expression in line with our observations in H7N9-infected patients who survived[30]. CD38^+^MHC-II^+^PD-1^+^CD8^+^ T cells were also detected in the primary response following infection with the mouse adapted strain, X31. Of note, increased disease severity resulting from infection with a high dose of X31 was not only characterized by excessive cytokine production, reminiscent of early hypercytokinemia observed during fatal H7N9 infection in humans[9], but also led to increased PD-1^+^ expression in CD38^+^MHC-II^+^CD8^+^ T cells. This is expected given that CD8^+^ T cell exhaustion, defined by PD-1^high^ expression, is a notable feature of disease conditions where excessive or prolonged inflammatory conditions are present, such as during acute and chronic infections caused by highly virulent IAV strains[10], HIV[31],[32], B type hepatitis (HBV)[33] as well as lymphocytic choriomeningitis virus (LCMV)[34].

Within the total primed CD38^+^MHC-II^+^ CD8^+^ T cell population in mice following influenza virus infection, a large proportion appeared to be tetramer-negative, which indicates recruitment of bystander CD8^+^ T cells at the site of infection. This was observed within populations of endogenous primary CD38^+^MHC-II^+^ CD8^+^ T cell populations (Fig.3) and during the secondary recall of OT-I cells (Fig.4). This could be due to inflammation-induced expansion not only at the site of infection but also in peripheral immune organs as numerous studies show that cytokines[35] such as type I IFN[36], IL-15[37], IL-12/IL-18[38] are capable to recruit non-virus-specific CD8^+^ T cells during different infectious diseases. It is unclear however what impact these bystander CD38^+^MHC-II^+^ CD8^+^ T cells have on the overall immune response against IAV infection. Similar findings have been noted by researchers who found bystander CD8^+^ T cells activated during the early phases of HIV-1 infection with CD38^+^MHC-II^+^ phenotype and specificity for cytomegalovirus, Epstein–Barr virus and influenza antigens[20]. Moreover, bystander CD8^+^ T cells and CD38^+^MHC-II^+^ CD8^+^ T cells can exhibit cytotoxic effects during acute Hepatitis A virus infection closely correlating with severe liver injury[39]. Thus, while not much is known about the function of bystander CD38^+^MHC-II^+^ CD8^+^ T cells during influenza infection, it is possible that their presence may be related to the hypercytokinemia state at site of infection and contribute to lung damage leading to severe disease outcomes[9].

In contrast to human CD8^+^ T cells, murine CD8^+^ T cells lack the transcription factor *CIITA*, a master regulator of *H2-Ab (MHC-II)* gene expression[25]. However, MHC-II is highly expressed on antigen-presenting cells (APCs) and our results are the first to demonstrate *in vivo* that CD38^+^CD8^+^ T cells can “snatch” MHC II, along with CD19 and possibly other cell surface markers from B cell populations. Trogocytosis of cell surface markers has been described for many immune cells in both mice and humans. For example, basophils can acquire MHC-II from dendritic cells to promote T cell differentiation to Th2 cells[40]. Similarly, T cells can acquire HLA-G from APCs changing their function as effector cells to regulatory cells[41]. Of relevance, CD38 expressed on B cells can establish lateral associations with various membrane proteins/complexes, including CD19 to help mediate intracellular signaling[42]. The interaction of both these markers thus paints a plausible scenario where CD19 is ‘snatched’ from B cells via interaction with CD38 on CD8^+^ T cells. Recent evidence also indicates that the exchange of markers between CD8^+^ T cells and B cells populations is bi-directional. Strittmatter-Keller *et al* in particular have reported a population of human B cells expressing CD8 despite the absence of CD8α or CD8β message, suggesting that this marker is acquired via cell-to-cell interactions with T cells[43].

What is to be gained by CD38^+^CD8^+^ T cells acquiring MHC II expression? Our adoptive transfer results demonstrate that MHC-II on murine effector CD38^+^CD8^+^ T cells plays a role in their optimal survival and expansion following secondary challenge. In fact, *in vitro* studies have shown that murine CD8^+^ T cells that have acquired MHC-II molecules from dendritic cells are able to stimulate CD4^+^ T cells in culture[44]. Given the importance of helper T cells in memory CD8^+^ T cell formation[45],[46] and in the recall of secondary responses[47],[48], it can be assumed that CD8^+^ T cells expressing MHC-II molecules can also present antigen for recognition by CD4^+^ T cells to stimulate the secretion of cytokines that promote effector function and differentiation into memory populations. This could therefore constitute an additional mechanism to amplify T-helper responses[49] but may also contribute to excessive responses or even tolerance[50], especially if immune-regulatory mechanisms are perturbed as reflected in dysfunctional innate immune responses observed during severe H7N9-infection[9]. Other implications can be gleaned from studies demonstrating snatching of peptide-MHC-I complexes (pMHC) by CD8^+^ T cells from APCs via cell-cell interactions[51],[52],[53]. Here, trogocytosis can lead to fratricide of T cells or exhaustion due to prolonged engagement of TCRs with “snatched” MHC-II-antigen complexes especially in the presence of high densities of pMHC[51]. These scenarios highlight the impact of trogocytosis of MHC-II molecules, enhancing interaction between CD4^+^ and CD8^+^ T cells to boost memory CD8^+^ T cell recall responses but in the event of a severe viral infection, this process may impair CD8^+^ T cell responses as observed in fatal H7N9 patients.

## Materials and Methods

### Ethics statement

Animal experiments were conducted according to the Australian National Health and Medical Research Council Code of Practice for the Care and Use of Animals for Scientific Purposes Guidelines for the housing and care of laboratory animals. Ethics was approved by the University of Melbourne Animal Ethics Experimentation Committee (1714108). Animal experiments involving H7N9 infection were approved by the Shanghai Public Health Clinical Center in China (2014-E032-01).

### Mice and viral infections

For primary infections, 6-8 week old male or female C57BL/6 mice were lightly anaesthetised with isofluorane and infected by intranasal (i.n.) instillation (30μl) with 10^2^ and 10^5^ PFU of A/HK/x31 (X31; H3N2). To examine secondary CD8^+^ T cell responses in mice following H7N9 infection, mice were first primed i.n. with 10^4^ EID_50_ of A/Chicken/Shanghai/F/98 (H9N2) virus, followed 60 days later with 10^6^ pfu of A/Shanghai/4664T/2013 (H7N9) virus.

To investigate mechanisms underlying MHC-II expression on murine CD8^+^ T cells, 10^4^ naïve OT-I cells were adoptively transferred into C57BL/6 mice or MHC-II−/− mice 24 hours prior to infection with X31-OVA. After 8 days, CD45.1^+^CD8^+^Vα2^+^ OT-I cells in the BAL, mediastinal lymph node and spleen were assessed. The experiment was repeated twice with 3-5 mice per group for each repeat.

To examine the recall ability of CD38^+^MHC-II^+^CD8^+^ T cells, 10^4^ naïve OT-I cells were adoptively transferred into C57BL/6 mice 24 hours prior to infection with X31-OVA. After 8 days, CD45.1^+^CD8^+^Vα2^+^ OT-I cells in the lymph node and spleen were four-way sorted based on their expression of CD38 and MHC-II. Each sorted population was adoptively transferred into separate naïve C57BL/6 recipient mice (5-10^3^ cells per mouse) and 30 days later infected with X31-OVA. The experiment was repeated twice with 3-5 mice per group for each repeat.

### Tissue sampling and cell preparation

Lungs, spleens, mediastinal lymph nodes (MLN) and bronchoalveolar lavage (BAL) were collected from mice at various time points after infection. Lungs were either homogenized and centrifuged to obtain clarified supernatants to assay for viral titres or enzymatically digested in collagenase III (Worthington Biochemical Corporation, USA; 1mg/ml) and DNase I (Sigma-Aldrich, Germany; 0.5mg/ml) before passing through cell sieves to obtain single cell suspensions for analysis. Where necessary, cell suspensions from tissues were incubated with 0.15M NH4Cl and 17 mM Tris- HCI at pH 7.2 for 5 mins at 37°C to lyse red blood cells.

### Measurement of lung viral load

Titres of infectious virus were determined by plaque assays on confluent Madin Darby canine kidney (MDCK) cell monolayers cultured in six-well plates. Cells were infected with lung homogenate supernatants at varying dilutions for 45 min at 37°C before the addition of Leibovitz L15 or MEM medium containing 0.9% agarose overlay containing Trypsin (Worthington Biochemical, NJ, USA), as previously described[54]. Plates were incubated at 37°C 5% CO2 for 3 days and virus-mediated cell lysis then counted as plaques on the cell layer and expressed as plaque forming units (PFU).

### Analysis of cytokine levels

Cytokines present in lung homogenate supernatants were measured using a BD CBA flex set (BD Bioscience) as per the manufacturer’s instructions[54]. Samples were analysed using a Becton Dickinson FACS Canto II flow cytometer. Data were analysed using FCAP Array software (Soft Flow Inc., Pecs, Hungary).

### Tetramer and antibody staining

MHC-I tetramer targeting the immunodominant epitope of the influenza nucleoprotein (DbNP366-374 – ASNENMETM, DbPA224-233 - SSLENFRAYV) was produced in-house and conjugated to streptavidin-APC/PE (Life Technologies Australia Pty Ltd) at a 1:250 dilution at room temperature for 1h. Cells were stained with combinations of fluorochrome-conjugated antibodies.PerCP-Cy5.5-CD3(#551163), PE-CD8(#561095), BV421-I-Ab(#562928), APC-CD44(#553133), FITC-CD44((#553133), BV711-CD38(#740697), PerCP-Cy5.5-CD8(#551162), APC/eF780-CD62L(#47-0621-82), APC-CD11c(#17011481), PE-Cy7-TCR-va2(#560624) from BD Biosciences, USA etc. PE-Cy7-CD38(#102718), BV785-PD-1(#329908), APC-Cy7-CD45.1(#110716), FITC-I-Ab(#116406), Percific Blue-I-Ab(#116422), FITC-CD19(#115506) from Biolegend, USA. AF700-CD3(#56003382) was purchased from Invitrogen, USA.

Live/Dead-aqua 525 were purchased from Invitrogen. Briefly, cell suspensions were stained with Live/Dead Aqua viability dye at room temperature for 10 mins followed by staining with tetramer for 15 mins and cell surface marker antibodies for 30 mins. Cells were fixed with 1% paraformaldehyde before analysis by flow cytometry. All antibody and tetramer staining was performed at 4°C and in the dark. Samples were subsequently acquired on a Becton Dickinson LSR Fortessa or Aria III flow cytometer and data analyzed by Flowjo Software (Tree Star Inc, USA).

### Statistical analyses

The comparison between absolute numbers of cells were analysed using two-way ANOVA analysis. Body weight between mice groups were compared by multi-T test. Absolute numbers of cells OT-I cells from different mice groups were compared using ordinary one-way ANOVA analysis.

## Supplemental material

Fig.S1 shows gating strategies on different subsets of CD8^+^ T cells and phenotypes.

Fig.S2 shows (a) early recruitment of CD38^+^MHC-II^+^ CD8^+^ T cells and influenza-specific CD8^+^ T-cell to BAL after severe influenza virus challenge and (b) gating strategies on CD38, MHC-II and PD-1 phenotype on different subsets of CD8^+^ T cells.

---

**Figure S1.**
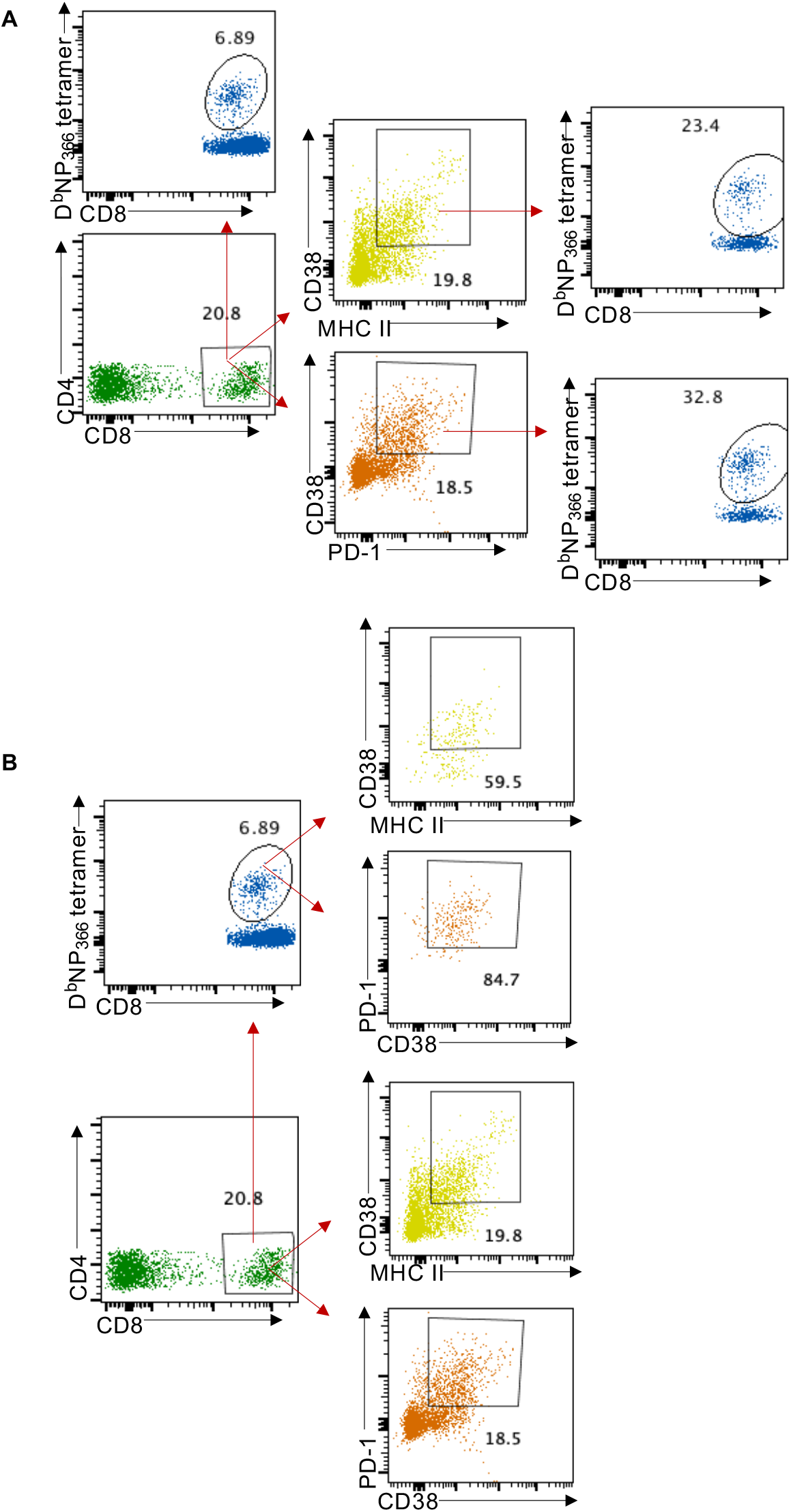
Gating strategy of D^b^NP_366_^+^CD8^+^ and CD38^+^MHC-II^+^CD8^+^ T cell populations. **(a)** Gating strategy to define D^b^NP_366_^+^CD8^+^ T cells from the pool of total CD8^+^ T cells, CD38^+^MHC-II CD8^+^ T cells or CD38^+^ PD-1^+^ CD8^+^ T cells. **(b)** Gating strategy to define CD38^+^MHC-II CD8^+^ and CD38^+^ PD-1^+^ CD8^+^ T cells from total CD8^+^ or D^b^NP_366_^+^CD8^+^ T cell populations.

**Figure S2.**
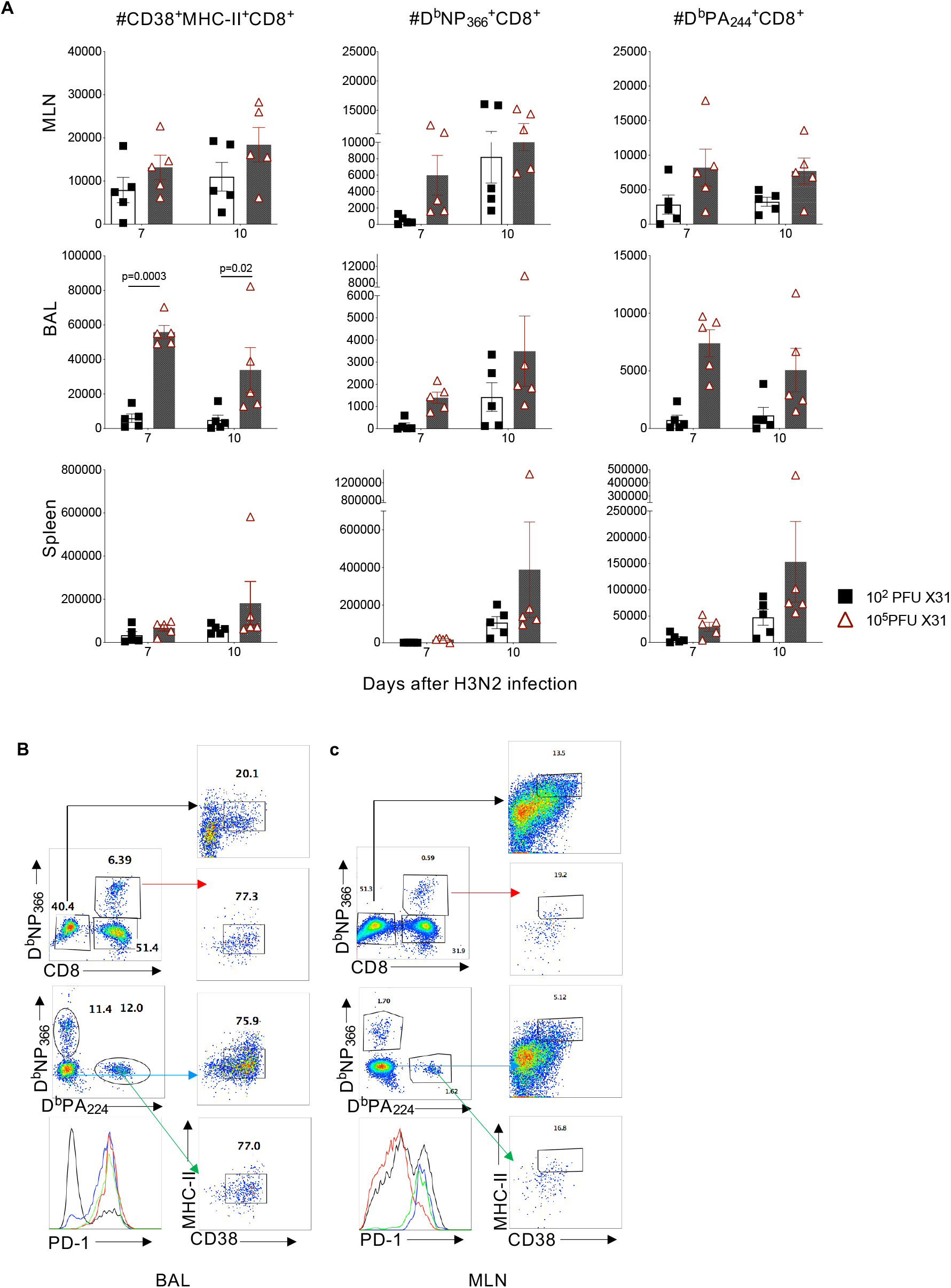
Early recruitment of tetramer-negative CD38^+^MHC-II^+^CD8^+^ T cells to the site infection with a high viral dose of H3N2. B6 mice were infected with HKx31 (H3N2) at either a high (10^5^ pfu) or low (10^2^ pfu) dose i.n.. At d7 and d10 after infection, BAL, spleen and MLN lymphocytes were stained with the D^b^NP366 and D^b^PA_224_ tetramers and anti-CD8 antibody, combined with a phenotyping panel for CD38, PD-1 and I-A^b^. (**a**) Numbers of CD38^+^MHC-II^+^CD8^+^ T-cells, D^b^NP_366_^+^CD8^+^ and D^b^PA ^+^CD8^+^ T cells are shown in MLN, BAL and spleen at d7 and d10 after infection (n=5 mice). (**b, c**) Representative FACS plots are shown for CD38, MHC-II and PD-1 expression on D^b^NP_366_^+^CD8^+^, D^b^PA ^+^CD8^+^ T cells and tetramer-negative CD8^+^ T cells.

## Acknowledgements

The authors thank Prof SJ Turner for insightful discussions. This work was supported by the Australian National Health and Medical Research Council (NHMRC) Program Grant (#1071916) and NHMRC Investigator Grant (#1173871) to KK. XJ was a recipient of XJ by China Scholarship Council-UoM Joint Scholarship.

Author contributions: XJ, ZW, BC, LL, LK and MK performed and analysed the experiments. JX, ZW, JQ, WH and KK designed experiments. XJ, BC and KK wrote the manuscript. All authors revised the manuscript. KK led the study.

## Competing Interests

### Disclosures

The authors declare no competing interests exist.

